# Pan-filovirus one-step reverse transcription-polymerase chain reaction screening assay

**DOI:** 10.1101/579458

**Authors:** Katharina Kopp, Ina Smith, Reuben Klein, Shawn Todd, Glenn A. Marsh, Alister C. Ward

## Abstract

Five species within the genera *Ebolavirus* and *Marburgvirus* of the family *Filoviridae* are known to cause severe hemorrhagic fever with high mortality rates in humans and non-human primates. Recent large outbreaks of Ebola virus disease in West Africa (2014 - 2016) and the Democratic Republic of the Congo (2018 - ongoing) have demonstrated the epidemic potential with devastating public health consequences. Several known and novel filovirus species have been found in bats in recent years. However, the role of each virus species in the disease ecology of human disease is still unclear. In particular, the transmission mechanism from potential animal hosts to humans is not known. Therefore, a simple, flexible, cost-effective screening tool for detecting the presence of any (putative) member of the filovirus family in animal samples is needed. In this study, a one-step conventional pan-filovirus RT-PCR assay was developed. The designed universal consensus primers of this screening test target two highly conserved regions of the nucleoprotein (NP) of all currently known filoviruses. The assay was capable of specific amplification of viral RNA of all six primate-pathogenic (human and non-human) filovirus species and resulted in 317 bp long RT-PCR products. This amplicon length renders the assay suitable for flexible application as conventional reverse transcription polymerase chain reaction (RT-PCR) as well as for future use as rapid real-time quantitative reverse transcription polymerase chain reaction (RT-qPCR).

## INTRODUCTION

Five species of the family *Filoviridae* are known to be pathogenic in humans. Acute infections with the four species *Zaire ebolavirus, Sudan ebolavirus, Tai Forest ebolavirus*, and *Bundibugyo ebolavirus* of the genus *Ebolavirus* and the single species *Marburg marburgvirus* of the genus *Marburgvirus* cause severe disease in human patients with often high case fatality rates (Peters & Khan, 1999, Feldmann & Geisbert, 2011, Emanuel et al., 2018). While increased bleeding tendencies are not necessarily present in all clinical cases infected with these filoviruses (WHO, 1978, Bwaka et al., 1999, Okware et al., 2002, MacNeil et al., 2010, Roddy et al., 2010, Schieffelin et al., 2014, Bah et al., 2015, McElroy et al., 2015, Rougeron et al., 2015, Miraglia et al., 2019), the diseases caused by them are classified under the syndromic cluster of viral hemorrhagic fevers (VHF). Large outbreaks of Ebola virus disease (EVD) caused by *Zaire ebolavirus* (28,652 cases in West Africa in 2014 - 2016, Bell et al., 2016) and *Sudan ebolavirus* (425 cases in Uganda in 2000-2001, Okware et al., 2002) as well as Marburg virus disease (MVD, 252 cases in Angola in 2004-2005, Towner et al., 2006) have demonstrated the epidemic potential in human populations.

Only in the most recent two outbreaks of EVD in the Democratic Republic of the Congo (Western region April - August 2018 and Eastern region 01 August 2018 - ongoing as of 21 February 2019) a national ethics committee has approved an experimental vaccine and four experimental therapeutics against the *Zaire ebolavirus* (WHO, 2018a, WHO, 2018b, WHO, 2018c, Ministere de la Sante de RDC, 2018, NIHCC, 2018). Persons at high risk, such as front-line health workers and contacts of patients, are vaccinated. Laboratory-confirmed patients are treated with emergency-approved drugs. Vaccines and treatments have also been developed for diseases caused by the other four human pathogenic filoviruses and some of them have been tested in animal models (Warren et al., 2014, Pittman et al., 2018). However, no officially licensed vaccine or therapy is available today.

This lack of medical countermeasures emphasizes the importance of prevention at the source. A single trans-species spillover event is commonly considered as starting point for consecutive human-to-human transmission chains in Ebola virus disease outbreaks, including the largest epidemic to date (Mylne et al., 2014, Pigott et al., 2014, Gire et al., 2014, Marí Saéz et al., 2015). While this sole spillover mechanism was also assumed for the largest Marburg virus disease epidemic in Angola in 2004 – 2005 (Towner et al., 2006, Pigott et al., 2015), multiple introductions from animal hosts were hypothesized for the second largest MVD outbreak in the Democratic Republic of the Congo in 1998 - 2000 (Bausch et al., 2006, Pigott et al., 2015).

However, there is a major gap in knowledge on the full range of natural reservoir host animals and possible vectors (Caron et al., 2018). The only filovirus species for which a bat species (Egyptian fruit bat, *Rousettus aegyptiacus*) has been confirmed as natural reservoir host by virus isolation and experimental infection is the *Marburg marburgvirus* (Towner et al., 2009, Amman et al., 2012, Amman et al., 2014, Paweska et al., 2012, Paweska et al., 2015, Amman et al., 2015). While this finding was the first evidence for the hypothesis that bats play an important role in filovirus disease ecology, it neither revealed the entire host spectrum of this particular filovirus species and the virus family as a whole nor the specific transmission mechanism from an animal host to a human patient. Characterization of the Měnglà virus in a fruit bat of the genus *Rousettus* in China, for which the creation of the novel filovirus genus *Dianlovirus* was suggested (Yang et al., 2019) is the most recent indicator that bats are natural reservoir hosts of filoviruses. The discovery of a new species member of the genus *Ebolavirus*, which was named *Bombali ebolavirus*, in an insectivorous bat species in Liberia (Goldstein et al., 2018) also supports this concept.

Non-human primates and duikers have been found to develop clinical signs of Ebola and Marburg virus disease and also succumb to it (Leroy et al., 2004, Rouquet et al., 2005, Wittmann et al., 2007, Gonzalez et al., 2007, Karesh et al., 2012). Therefore, hunting of sick wildlife and especially collecting dead animals of these species can be assumed as one source of infection for humans. However, large-scale screening of a wide range of animal species ranging from arthropods as potential vectors to vertebrates as reservoir and transmission hosts of filoviruses is necessary in order to cover all possible spillover interfaces with humans.

Several assays employing conventional (RT-PCR) and real-time reverse transcription polymerase chain reaction (RT-qPCR) have been developed for the molecular detection of filoviruses, as reviewed by Clark et al., 2018. Most of these tests were designed for rapid and sensitive laboratory diagnosis and confirmation of clinical disease caused by a previously identified filovirus species. As the majority of these nucleic acid detection assays target distinct sequences of certain filovirus species, they are not suitable for screening a broad range of different virus species in one assay, as would be required for ecological investigations.

With the purpose of reducing the complexity, time and costs of such screenings as well as to facilitate the discovery of novel filoviruses, we aimed for the development of a single molecular assay which can detect all known and possibly novel members of that virus family. As proof of principle for such an approach in screening potential natural hosts for all known and novel filoviruses, it was shown that the developed one-step RT-PCR assay was capable of specific amplification of viral RNA of isolates of *Marburg marburgvirus* and five species of the genus *Ebolavirus* in a universal optimized protocol.

## MATERIALS AND METHODS

### RNA extraction

For assay evaluation, viral RNA from Vero E6 culture supernatants of filoviruses, listed in Table 4, were extracted using the QIAamp Viral RNA Mini Kit (catalog-no. 52906, Quiagen, Venlo, the Netherlands), as recommended by the manufacturer. Since this treatment renders the sample of negative-sense viral RNA non-infectious, samples were dunked out from the biosafety level 4 (BSL-4) laboratory suite of the Commonwealth Scientific and Industrial Research Organisation (CSIRO) Health and Biosecurity, Australian Animal Health Laboratory, Geelong, Australia, aliquoted in 20 μl, and stored at −80°C for further investigations.

### Oligonucleotide design

All coding sequences (CDS) of the family *Filoviridae* (GenBank taxonomic identity 11266), as available on 17 April 2015, were downloaded from the GenBank database (https://www.ncbi.nlm.nih.gov/nuccore/?term=txid11266[Organism:exp]). A parsing script using the term “gene=NP” filtered the respective FASTA file, in order to contain only the nucleoprotein (NP) CDS.

The resulting NP nucleotide sequences were aligned by the standalone version of the multiple sequence alignment (MSA) tool Clustal Omega (https://pkgs.org/download/clustal-omega, version 1.2.4, the web server version can only process input files < 4 MB) and visualized by the MSA editor and analysis workbench Jalview (Waterhouse et al., 2009). A target region covering a total length of 317 bases of filovirus NP sequences and including two highly conserved areas of 28 nucleotide length at each end was identified by visual inspection of the MSA.

Sequences containing gaps or not clearly identified bases (“N”) within the two highly conserved ends of the target region were deleted. Unique sequences of this target area were kept for the design of the universal consensus primers, while identical sequences within this region were removed by using Jalview’s “remove redundancy” option. Two sense and two antisense universal consensus primer sequences were designed and chosen for specificity for the family *Filoviridae* by using the tool Primer-BLAST (https://www.ncbi.nlm.nih.gov/tools/primer-blast/, Ye et al., 2012) and are listed in Table 1.

**Table 1:**
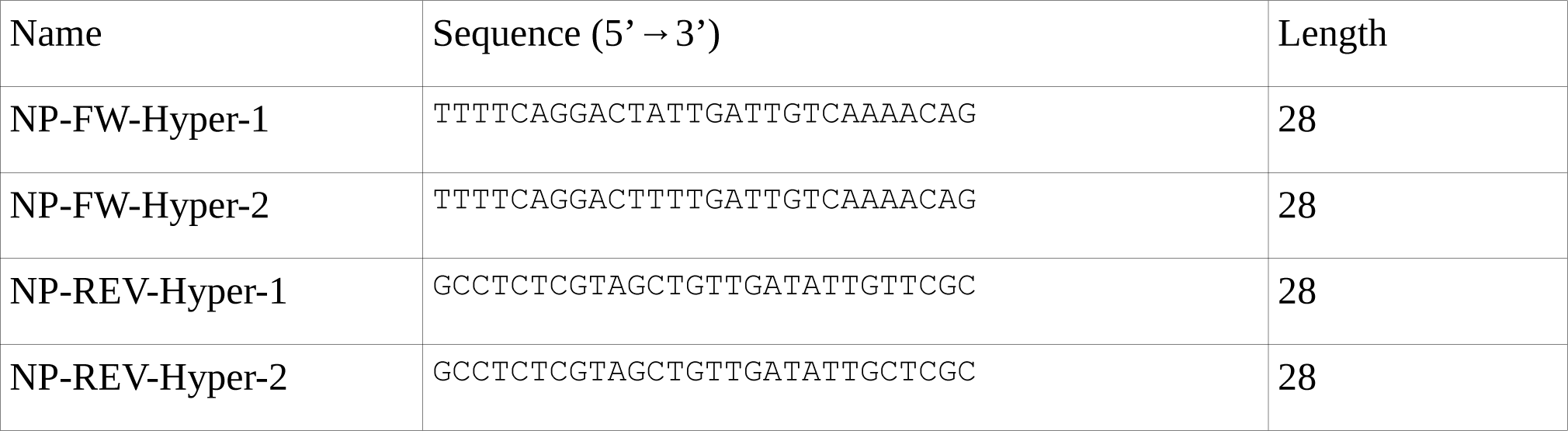
Universal consensus primer set targeting the nucleoprotein region of filoviruses.

### Conditions of the one-step pan-filovirus RT-PCR assay

The SuperScript™ III One-Step RT-PCR System with Platinum™ Taq DNA Polymerase (catalog- no. 12574-026, Invitrogen™, Thermo Fisher Scientific, Waltham, Massachusetts) was used in the pan-filovirus RT-PCR screening assay. The 20-μl assay contained 10 μl of 2X reaction mix provided with the kit (including the basic level of 1.6 mM of MgSO_4_), 0.4 μl of each of the four 10μM primers (Hyper-NP-FW-1, −FW-2, −REV-1, −REV-2), 0.8 μl of SuperScript™ III RT/Platinum™ Taq enzyme mix, and 3.6 μl of nuclease-free water (NFW). Finally, for the screening of samples 4 μl of extracted RNA was added, for no template controls (NTC) 4 μl of NFW, or for positive controls 4 μl of *in vitro* transcribed antisense ssRNA. All reactions were conducted in duplicate.

### Cycling profile of the one-step pan-filovirus RT-PCR assay

The RT-PCR assay involved the following steps: reverse transcription at 50°C for 15 min; reverse transcriptase inactivation and hot-start activation at 94°C for 2 min; 10 touch-down precycles with 4°C for 15 s, 68°C for 30 s with a temperature decrease of 1°C per cycle (starting at cycle 2 out of 10), and 68°C for 1 min; 40 cycles with 94°C for 15 s, 60°C for 30 s, and 68°C for 1 min; and final extension at 68°C for 5 min.

### Results visualization of the one-step pan-filovirus RT-PCR assay

The 1% agarose (UltraPure™ Agarose, catalog-no. 16500500, Invitrogen™, Thermo Fisher Scientific Waltham, Massachusetts) gel was cast using 1x SYBR™ Safe DNA Gel Stain (catalog- no. S33102, Invitrogen™, Thermo Fisher Scientific Waltham, Massachusetts) and 1× Tris-EDTA buffer (UltraPure™ DNA Typing Grade™ 50X TAE Buffer, catalog-no. 24710030, Invitrogen™, Thermo Fisher Scientific Waltham, Massachusetts). Amplicons generated by the one-step pan-filovirus RT-PCR assay were resolved by gel electrophoresis running at 100 volts for an average of 40 minutes, visualized and captured using the Safe Imager™ 2.0 Blue-Light Transilluminator system (catalog-no. G6600EU, Invitrogen™, Thermo Fisher Scientific Waltham, Massachusetts). The 1 Kb Plus DNA Ladder (catalog-no. 10787018, Invitrogen™, Thermo Fisher Scientific Waltham, Massachusetts) was used as a marker for estimating the approximate sequence length of PCR products.

### Product sequencing of the one-step pan-filovirus RT-PCR assay

Gel bands of PCR products of samples which were found to have a size approximately corresponding to the 317 bp amplicon, as expected for true positives of all known filovirus sequences (as of 17 April 2015 in the GenBank database, https://www.ncbi.nlm.nih.gov/nuccore/), were excised, purified and prepared for capillary sequencing using the BigDye™ Terminator v3.1 Cycle Sequencing Kit (catalog-no. 4337455, Applied Biosystems™, Thermo Fisher Scientific Waltham, Massachusetts). For each resulting amplicon sequence a search over the entire “nucleotide collection (nr/nt)” database by using the “blastn suite” https://blast.ncbi.nlm.nih.gov/Blast.cgi?PROGRAM=blastn) of the BLAST algorithm (Altschul et al., 1990) was conducted.

### *In vitro* transcription of the antisense ssRNA positive control

Antisense ssRNA molecules of the full-length nucleoprotein (NP) coding sequence (GenBank ID AF522874, position 464-2683) of the *Reston ebolavirus* strain Pennsylvania (Groseth et al., 2002) were *in vitro* transcribed as virus-specific positive controls. While not used as positive controls, also sense ssRNA molecules of the full-length NP CDS of this *Reston ebolavirus* strain were *in vitro* transcribed to evaluate if the developed one-step pan-filovirus RT-PCR assay can specifically amplify ssRNA molecules of both directions. The ∼2.2 kb diagnostic target was amplified from a pCAAGGS vector (“pCAGGSEboVRestonNP”) containing the entire NP region by using the primer pair ERp-NP-T7-antisense-FW (5 -TAATACGACTCACTATAGGGTTACTGATGGTGCT-GCAAGATTGC-3’) and Erp-NP-T7-antisense-REV (5’-GTAGGAAAAAGAAGAAGGCATGA-ACAT-3’) or the primer pair ERp-NP-T7-sense-FW (5 -TAATACGACTCACTATAGGGATG-TTCATGCCTTCTTCTTTTTCCTAC-3’) and ERp-NP-T7-sense-REV (5 -TTACTGATGGTGC-TGCAAGATTGC-3’). Correct length of the PCR products was verified by gel electrophoresis (Figure 1), the corresponding bands excised from the agarose 1% gel, and purified using the Wizard® SV Gel and PCR Clean-Up System (catalog-no. A9282 Madison, Wisconsin, USA).

**Figure 1:**
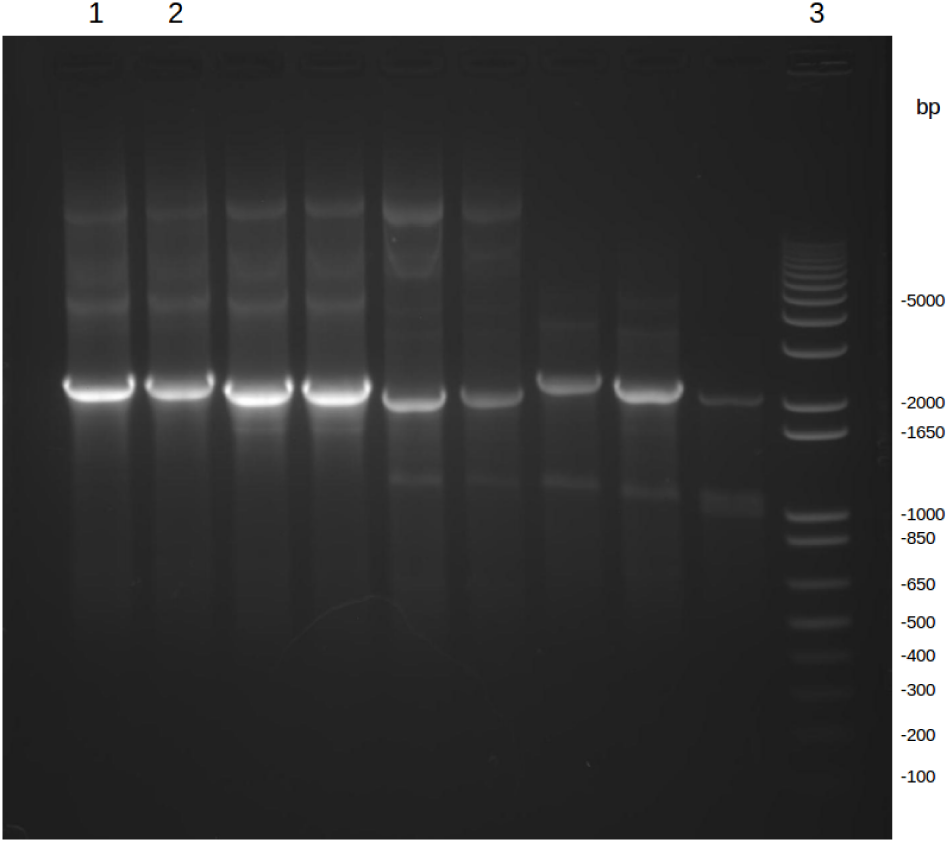
Gel electrophoresis of products of PCR with NP gene- and species-specific primers (lane 1: Erp-NP-T7-antisense-FW and Erp-NP-T7-antisense-REV, lane2: Erp-NP-T7-sense-FW and Erp-NP-T7-sense-REV) from circular NP plasmid DNAs. Lane 3: 1kb plus DNA ladder. Non-marked lanes from different experiments (Kopp, 2015, unpublished).

The purified PCR products were used as templates for *in vitro* transcription of antisense and sense ssRNA by using the HiScribe™ T7 In Vitro Transcription Kit (NEB #E2030S, New England BioLabs Inc, Ipswich, Massachusetts, USA). Finally, the *in vitro* transcribed antisense and sense ssRNA were purified by using the RNeasy® Mini Kit (catalog-no. 74106, Quiagen, Venlo, the Netherlands) including an on-column DNase I digestion step using the RNase-Free DNase Set (catalog-no. 79254, Quiagen, Venlo, the Netherlands). Before being employed as positive controls in the RT-PCR screening of samples for possible detection of filovirus RNA, the concentration of the *in vitro* transcribed antisense ssRNA control was determined by using the Qubit™ RNA BR Assay Kit (catalog-no. Q10210, Invitrogen™, Thermo Fisher Scientific Waltham, Massachusetts) and Qubit® Fluorometer 2.0 as ∼218 ng/μl.

The presence of the target region of the pan-filovirus RT-PCR assay within the full-length NP-coding sequence of the positive control antisense ssRNA as well as of the sense ssRNA molecule was verified by a *Reston ebolavirus*-specific RT-PCR employing the primers NP-FW-2 (5’-TTTTCAGGACTCCTAATTGTCAAAACCG-3’) and NP-REV-12 (5’-GCCTCTCTGAGCTGC-TGATACTGCTCAC-3’) (Figure 2). In addition, the capability of the developed one-step pan-filovirus RT-PCR assay (using the primers listed in Table 1) to detect sense and antisense ssRNA was evaluated (Figure 3). The absence of any target region DNA (which might have remained from previous steps in the *in vitro* RNA-transcription procedure) was verified by non-amplification in a no reverse transcriptase control PCR (minus-RT) reaction using the antisense ssRNA as a template and either the *Reston ebolavirus*-specific primer pair (Figure 2) or the universal consensus primer pair (Figure 3). Plasmid “pCAGGSEboVRestonNP” served as a positive control template in both minus-RT PCR reactions (Figure 2 and 3).

**Figure 2:**
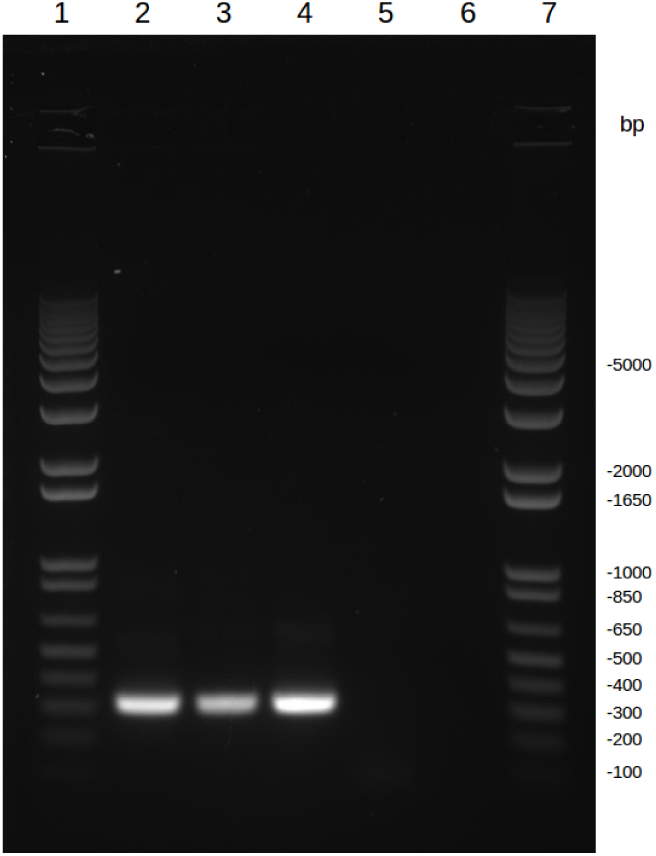
Detection of *in vitro* transcribed ssRNA by conventional one-step RT-PCR using the *Reston ebolavirus* species-specific primers NP-FW-2 and NP-REV-12 and as a template *in vitro* transcribed antisense ssRNA (lane2) or sense ssRNA (lane 3). Minus-RT PCR reaction using the species-specific primer pair NP-FW-2 and NP-REV-12 and as a template plasmid “pCAGGSEboVRestonNP” (lane 4) or antisense ssRNA (lane 5). No template control (NTC, lane 6). Marker: 1kb plus DNA ladder (lane 1 and 7).

**Figure 3:**
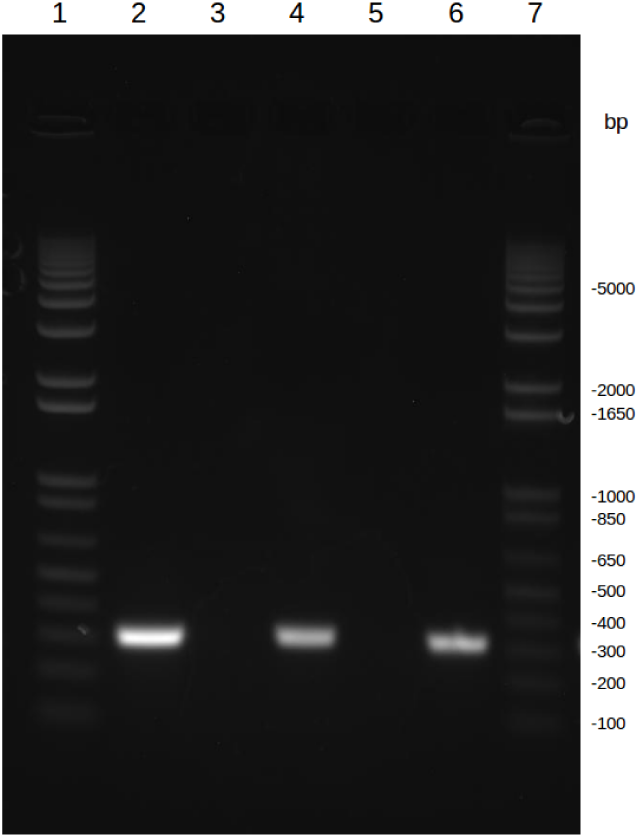
Detection of *in vitro* transcribed ssRNA by conventional one-step RT-PCR using the universal consensus primers NP-FW-Hyper-1, NP-FW-Hyper-2, NP-REV-Hyper-1, and NP-REV-Hyper-2 (Table 1) and as a template *in vitro* transcribed antisense ssRNA (lane 4) or sense ssRNA (lane 6). Minus-RT PCR reaction using the universal consensus primers and as a template plasmid “pCAGGSEboVRestonNP” (lane 2) or antisense ssRNA (lane 3). No template control (NTC, lane 5). Marker: 1kb plus DNA ladder (lane 1 and 7).

## RESULTS

### Design of primers for the one-step pan-filovirus RT-PCR assay

The universal consensus primer set of the one-step pan-filovirus RT-PCR assay was developed according to all nucleotide sequences of the nucleoprotein gene of filoviruses as available on 17 April 2015 in the GenBank database (223 sequences). However, because of the large numbers of sequences determined from the *Zaire ebolavirus* epidemic in West Africa in 2014 - 2016 as well as the discovery of novel putative members of the family *Filoviridae*, recently, an updated (as of 8 Jan 2019) multiple sequence alignment (MSA) was created as described for the original MSA under Material and Methods. First, the new MSA consisted of 2,219 GenBank sequences spanning the entire target region of the assay. After removal of all sequences which showed ambiguous base calls (“N”) or caused gaps in the MSA within the two 28 bases long regions corresponding to the forward and reverse primers, 2,204 and 2,202 sequences remained, respectively. Unique sequences of the target area of the sense and antisense universal consensus primer regions were identified by using Jalview’s “remove redundancy” option.

The target regions of the two sense (NP-FW-Hyper-1 and NP-FW-Hyper-2) and two antisense (NP-REV-Hyper-1 and NP-REV-Hyper-2) universal consensus primers correspond to the genome positions 1190-1217 and 1479-1506 (in reverse complement orientation of the antisense primers), respectively, of the Ebola virus isolate “Mayinga, Zaire, 1976” (species: *Zaire ebolavirus*, GenBank ID AF086833.2). Table 2 and 3 display the MSAs of all unique sequence variations found for the target regions of the forward and reverse universal consensus primers.

**Table 2:**
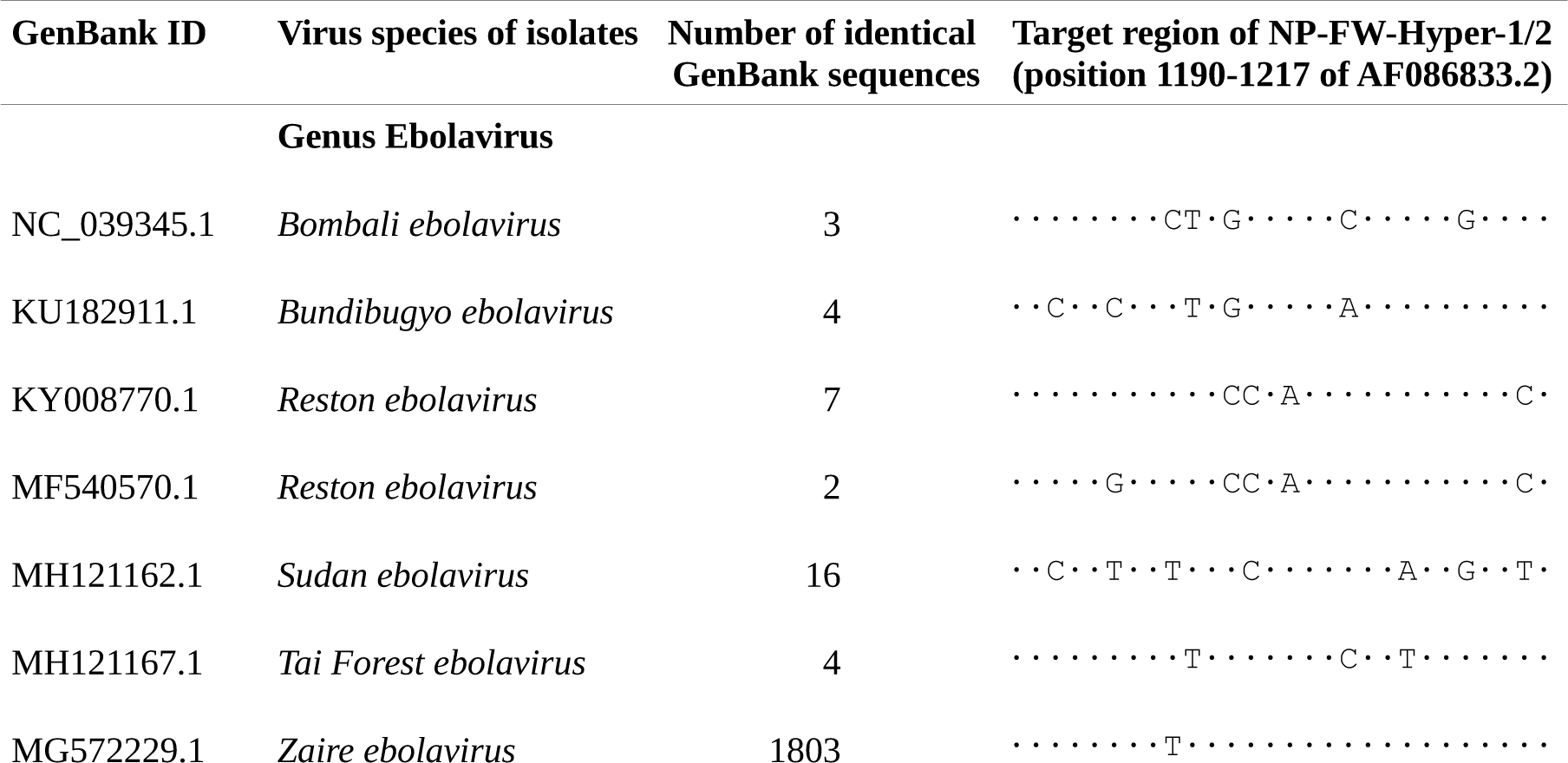
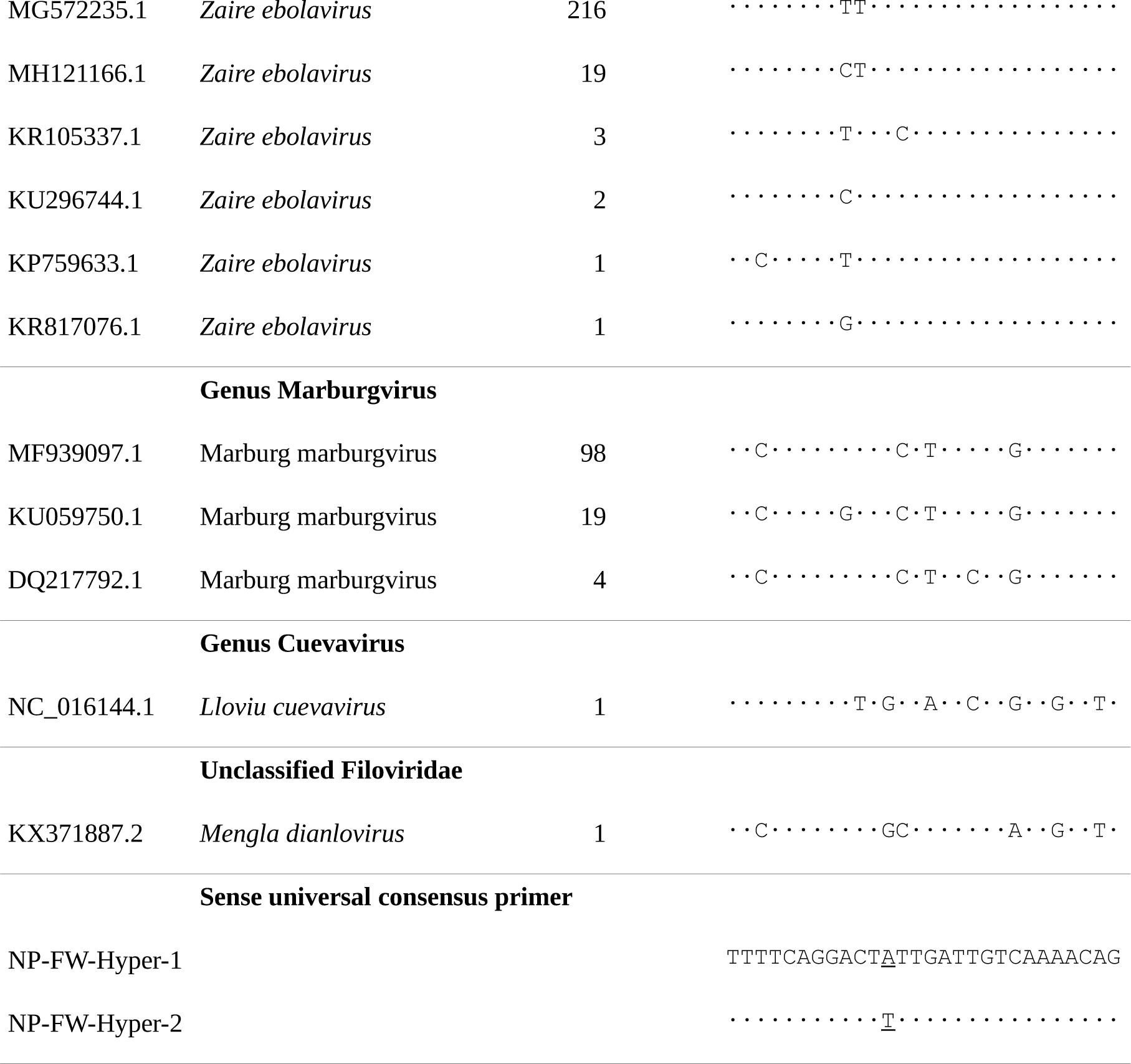
Target region of the designed universal consensus forward primers (NP-FW-Hyper-1 and NP-FW-Hyper-2) of the one-step pan-filovirus RT-PCR assay. All sequences of the target region available from the GenBank database were aligned and processed as described in the section Material and Methods. Every type of unique sequence variation in this region is represented by a GenBank accession number (column 1) and the filovirus species assigned to all GenBank entries of the identical sequence (column 2). The number of GenBank accessions (column 3) found for each unique sequence variant (column 4) is listed. Dots indicate sequence positions matching with that of the two sense universal consensus primer variants. Mismatches are represented by the base in the unique sequence variant. The two different positions of the two primers are underlined.

**Table 3:**
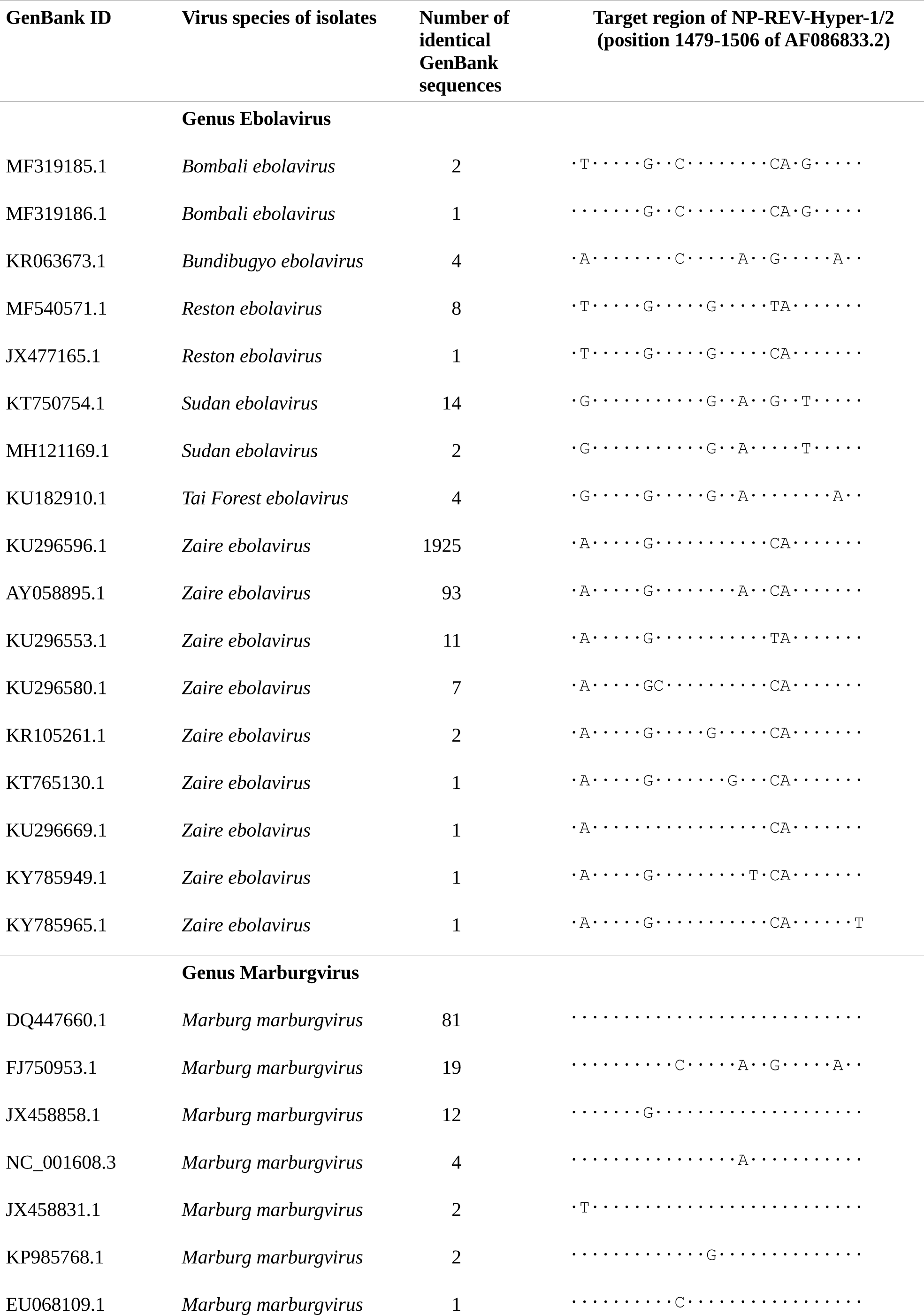
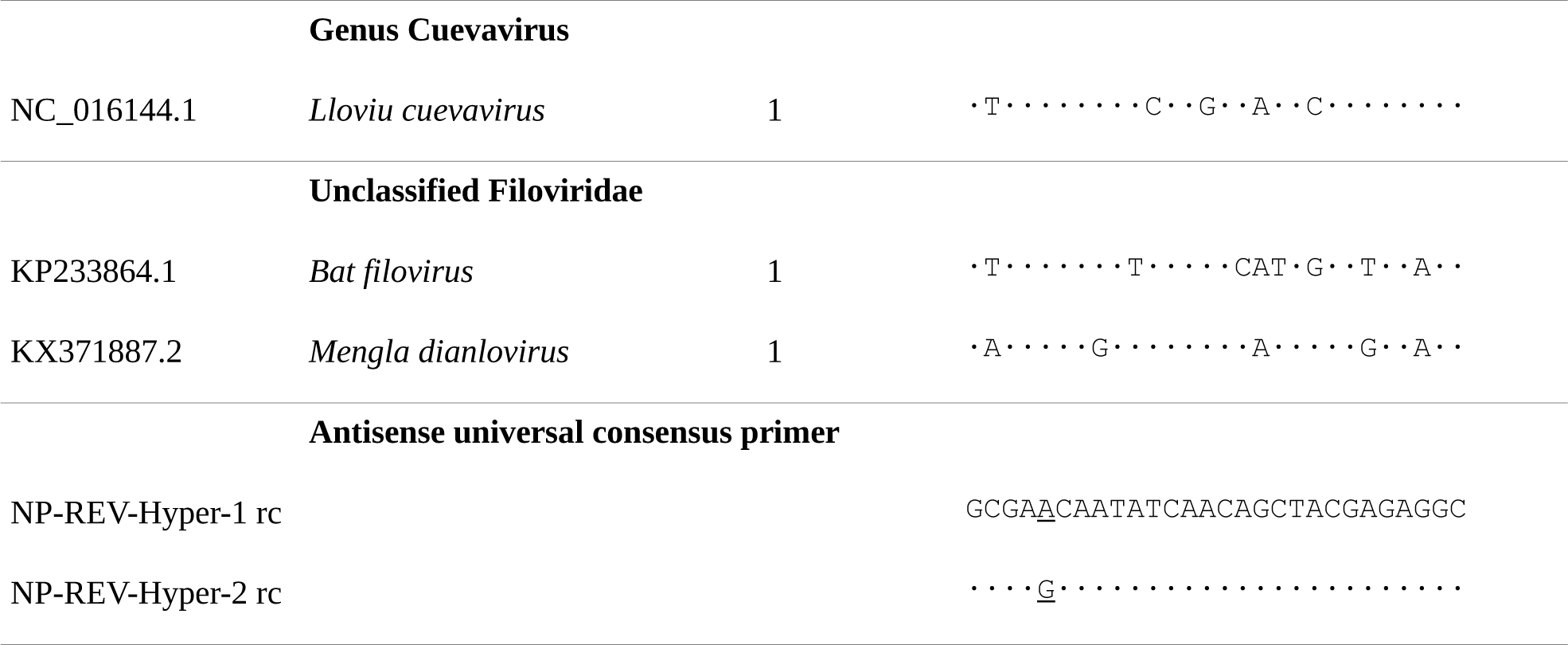
Target region of the designed universal consensus reverse primers (NP-REV-Hyper-1 and NP-REV-Hyper-2, here shown as reverse complement) of the one-step pan-filovirus RT-PCR assay. Legend and details as described in the caption of Table 2.

Each distinct sequence is represented by one GenBank accession. All identical sequences of each distinct variation were found to belong to isolates of the same filovirus species. For the majority of filovirus species (within the genus *Ebolavirus*: *Bombali ebolavirus, Bundibugyo ebolavirus, Reston ebolavirus, Sudan ebolavirus*, and *Tai Forest ebolavirus*; within the genus *Cuevavirus*: *Lloviu cuevavirus*), one or two distinct sequences were determined in the target regions of the sense and antisense primers. Two unclassified filoviruses which have recently been discovered in Chinese fruit bat species, the *Bat filovirus* (GenBank ID KP233864, He et al., 2015) and the *Mengla dianlovirus* (GenBank ID KX371887, Yang et al., 2017), were also aligned. However, the partial nucleoprotein CDS available for the *Bat filovirus* does not cover the target region of the sense primer.

More than two unique sequence variations were only found in the forward and reverse primer target regions of the two filovirus species *Zaire ebolavirus* and *Marburg marburgvirus*. However, the majority of GenBank entries were identical with only one particular sequence variation. These were 1,803 identical entries out of a total of 2,045 GenBank accessions (88.2%) in the forward primer target region and 1,925 identical entries out of a total of 2,042 GenBank accessions in the reverse primer target region (94.3%) for *Zaire ebolavirus*. The respective figures found for *Marburg marburgvirus* were 98 identical entries out of a total of 121 GenBank accessions (81.0%) and 81 identical entries out of a total of 121 GenBank accessions (67.0%).

### Optimization of reaction and cycling conditions of the one-step pan-filovirus RT-PCR assay

An optimal final primer concentration of 0.4 μM was determined for each of the four universal consensus primers (Hyper-NP-FW-1, Hyper-NP-FW-2, Hyper-NP-REV-1, and Hyper-NP-REV-2) in a reaction by titration experiments, in comparison to the sensitivity of the assay when the *Reston ebolavirus*-specific primer pair was used at the (manufacturer-recommended) constant concentration of 0.2 μM. For this purpose, five different concentrations (0.15 μM, 0.2 μM, 0.3 μM, 0.4 μM, and 0.5 μM) of each of the two sense primers (Hyper-NP-FW-1 and Hyper-NP-FW-2) were simultaneously tested versus these five distinct concentrations of each of the two antisense primers (Hyper-NP-REV-1 and Hyper-NP-REV-2). These 5 × 5 = 25 reaction set-ups were examined in triplicate in detecting the *in vitro* transcribed positive control ssRNA in a 10-fold dilution series from 872 ng to 8.72 fg per reaction.

None of the reactions containing the 25 different combinations of concentrations of the universal consensus primer quadruplet produced a visible amplicon band in the following gel electrophoresis at a template concentration of 8.72 fg, which corresponds to ∼7,360 copies of the ∼2.2 kb ssRNA positive control template. Only one out of the three identical reactions which employed the 200 nM concentration of the virus-specific primer pair (NP-FW-2 and NP-REV-12) were found to give a specific amplicon signal in the gel electrophoresis at that template concentration.

The estimated limit of detection (LOD) was defined as the lowest ssRNA amount of the dilution series for which positive detection in all three identical reactions could be determined. For the assay using the virus-specific primer pair, the LOD was found to be 87.2 fg (or ∼73,600 copies) of the positive control ssRNA per reaction. The LOD value for the virus-specific primer pair assay was only reached by the reactions containing the 400 nM concentration of each of the four universal consensus primers, which was, therefore, considered as optimal final primer concentration of the pan-filovirus assay.

Analogously, the highest annealing temperature (in the 40 cycles after the 10 touch-down precycles) which was found to achieve the same sensitivity for the reactions using the four universal consensus primers (Hyper-NP-FW-1, Hyper-NP-FW-2, Hyper-NP-REV-1, and Hyper-NP-REV-2) and for the reactions using the positive-control-specific primer pair (NP-FW-2 and NP-REV-12) was 60°C.

### Specific amplification of viral RNA of isolates of *Marburg marburgvirus* and five species of the genus *Ebola virus* by the one-step pan-filovirus RT-PCR assay

Using the above-described reaction conditions and cycling profile, the developed pan-filovirus RT-PCR assay detected viral RNA extracted from Vero E6 cell culture virus isolates shown in Table 4. As the end-point RT-PCR pan-filovirus assay developed in this study, served as a first proof-of-principle application aiming for the future development of corresponding reverse transcription quantitative real-time PCR (RT-qPCR) pan-filovirus assays, specific amplification of viral RNA was conducted only with unquantified RNA extracts from virus cell culture supernatants (undiluted, and ten-fold dilutions from 10^−1^ to 10^−6^).

**Table 4:** Virus isolates for which the developed pan-filovirus RT-PCR assay detected the presence of viral RNA.

| Genus | Species | Virus isolate | GenBank ID |
| --- | --- | --- | --- |
| *Ebolavirus* | *Bundibugyo ebolavirus* | Bundibugyo 2007 | FJ217161 |
|  | *Tai forest ebolavirus* | Cote d'Ivoire 1994 | FJ217162 |
|  | *Reston ebolavirus* | Pennsylvania 1989 | AF522874 |
|  |  | Philippines 2009 | JX477165 |
|  | *Sudan ebolavirus* | Boniface 1976 | FJ968794 |
|  |  | Gulu 2000 | AY729654 |
|  | *Zaire ebolavirus* | Mayinga 1976 | AF086833 |
|  |  | Kikwit 1995 | JQ352763 |
|  |  | Makona 2014 | KR781609 |
| *Marburgvirus* | *Marburg marburgvirus* | Musoke 1980 | DQ217792 |
|  |  | Ravn 1987 | DQ447649 |
|  |  | Angola 2005 | DQ447660 |

RNA extracts of all virus isolates and their dilutions down to 10^−4^ showed specific amplification in gel electrophoresis analysis for all virus species examined (Table 4). As an example, Figure 4 displays the detection of viral RNA from a virus isolate of the Angola 2005 strain of the *Marburg marburgvirus* (GenBank ID Q447660) by the one-step pan-filovirus RT-PCR assay. The specific nucleotide sequence of the 317 bp PCR product of each virus isolate, which had been purified from the agarose gel, was verified by Sanger sequencing and BLAST analysis. Extracted viral RNA of Hendra virus, Nipah virus, Measles virus, Rabies virus, Australian bat lyssavirus, and Borna virus were provided by high-containment laboratory groups of CSIRO Health and Biosecurity, Australian Animal Health Laboratory, Geelong, Australia and did not show specific amplification in the developed pan-filovirus RT-PCR assay.

**Figure 4:**
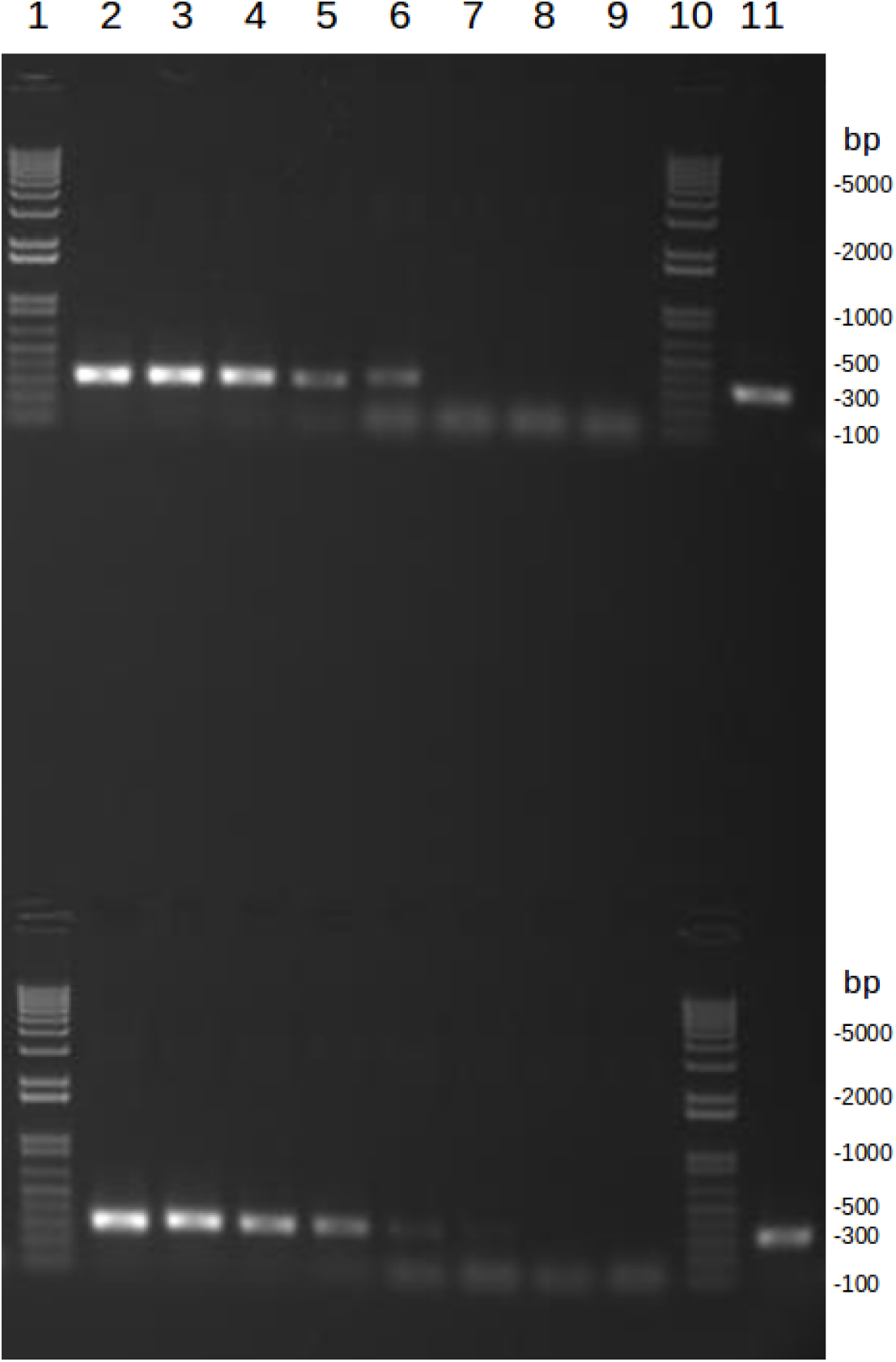
Detection of viral RNA extracted from a virus isolate of the Angola 2005 strain of the *Marburg marburgvirus* (GenBank ID Q447660) cell culture supernatant by conventional one-step RT-PCR using the universal consensus primers NP-FW-Hyper-1, NP-FW-Hyper-2, NP-REV-Hyper-1, and NP-REV-Hyper-2 (Table 1). Undiluted viral RNA extract (lane 2), dilutions 10^−1^ to 10^−6^ (lane 2 to 8), no template control (NTC, lane 9), and positive control using ∼8.72 pg of *in vitro* transcribed antisense ssRNA NP of *Reston ebolavirus* (lane 11). All reactions in duplicate. Marker: 1kb plus DNA ladder (lane 1 and 10).

## DISCUSSION

The one-step pan-filovirus RT-PCR assay developed in this study provides a robust and broad first screening tool in determining if an animal sample contains viral RNA of a known or unidentified putative member of the family *Filoviridae*. The universal consensus primer pairs of this RT-PCR target two highly conserved regions within the nucleoprotein gene. According to this primer design, the multiple alignment analysis (Table 2 and 3) indicates that specific amplicons could be expected not only from the 12 tested virus isolates (Table 4) but also when viral RNA of most, if not all, currently known members of the family *Filoviridae* are used as a template. In line with these bioinformatics findings, the developed RT-PCR assay produced specific amplicons of 317 bp length of the nucleoprotein coding region of the viral genomes, when used for testing viral RNA extracts of 12 distinct virus isolates spanning six filovirus species (Table 4).

Therefore, the products of these RT-PCR reactions could serve as appropriate sequencing templates in attempts to identify the filovirus species or strain in a sample. This is important, as for several distinct virus species and lineages of the genus *Ebolavirus* and *Marburgvirus* more specific and sensitive tests, such as reverse transcription quantitative real-time PCR (RT-qPCR) assays are available (Gibb et al., 2001, Drosten et al., 2002, Weidmann et al., 2004, Towner et al., 2004, Trombley et al., 2010, U.S. FDA, 2014, Pinsky et al., 2015, Southern et al., 2015, WHO, 2015, TIB MolBiol & Roche, n.d., Cnops et al., 2016, Rieger et al., 2016, Semper et al., 2016, Loftis et al., 2017, Pettitt et al., 2017). Overall, with hundreds of viral genome copies or less, the limit of detection (LOD) of each of these real-time RT-qPCR tests (Clark et al., 2018) is around two orders of magnitude lower than that (∼73,600 copies) found for the end-point RT-PCR developed in this study. The determined sequence of the amplicons could also inform the design of exactly matching primers (and probes, depending on the applied assay chemistry) for the development of more specific and sensitive follow-up nucleic acid detection assays.

Diagnosis and confirmation to investigate clinical samples in a human epidemic caused by an identified single filovirus and ecological screening of potential animal hosts for any member of this virus family require different methodological approaches. Several established real-time RT-PCR (RT-qPCR) systems are considered as “gold standard” in diagnosing filovirus infections in human patients. They, therefore, have been approved by the U.S. Food and Drug Administration (FDA) as methods of choice, in ongoing outbreak situations, mainly when the species or strain of the causative agent is known. As all of these methods target specific regions of one or more filovirus species whose nucleotide sequences had been determined at the time of the assays’ development, they are expected to be capable of reliably detecting only certain filovirus species.

As shown in the outbreak of the then unknown *Bundibugyo ebolavirus* in Western Uganda in 2007, using RT-(q)PCR assays, originally developed for different species or lineages of the virus family *Filoviridae*, can lead to 100% false-negative results (Towner et al., 2008). Only after virus isolation from patients and consecutive whole-genome sequencing a highly sensitive and specific RT-qPCR test was developed (Towner et al., 2008) and successfully applied in the control of the epidemic by correct patient identification, quarantine, and supportive treatment. The described management of this outbreak probably considerably contributed to its limited extent in duration and space compared with other previous filovirus epidemics but might have been implemented just in time. The initial complete failure of tests based on certain filovirus species in this case, demonstrated that even having a panel of highly specific molecular detection methods at hand might be insufficient in dealing with novel and emerging pathogens promptly.

However, as experienced in the largest Ebola virus disease epidemic in West Africa in 2014 - 16 (Marí Saéz et al., 2015), spillovers from a yet not unambiguously identified reservoir host into the human population have always started in a remote region of peri-equatorial Africa and must ideally be curbed at their source of origin. As witnessed in the warning example of the *Bundibugyo ebolavirus* outbreak in 2007 (Towner et al., 2008), developing an appropriate molecular detection method requires virus isolation and full-length sequencing of at least the RT-qPCR’s target region. These prerequisites demand sophisticated and high-containment (BSL-4) laboratory facilities and trained personnel, which might not be fast available, especially in resource-limited and remote areas.

While full-length genome sequencing and isolation from tissue culture are mandatory as final confirmation procedures for correct taxonomic classification of a potential filovirus and facilitating further research on diagnostics, treatment, and prevention, these rather complicated methods are not fast and robust enough for filovirus identification and discovery in the field. A broad range of detection capability at virus family level is needed for the first line of defense diagnostics in remote not yet identified outbreaks as well as in the search for the potential reservoir and transmission hosts. To close this gap, we developed a one-step RT-PCR assay which facilitates the screening of samples for the entire filovirus family in a simple, cost-effective “one-fits-all” protocol.

Three assays, two conventional RT-PCR tests (Sanchez et al., 1999, Zhai et al., 2007) and a real-time RT-qPCR test (Panning et al., 2007, partially using primers from Sanchez et al., 1999) also attempted to achieve this aim by targeting the L gene, which encodes the RNA-dependent RNA polymerase of filoviruses (Feldmann et al., 1993). However, the design of the consensus primer sets of all three assays had been based on the L gene sequences of all known filovirus species before the discovery of the Bundibugyo ebolavirus in 2007. However, a following modified version of the real-time RT-qPCR, initially developed by Panning et al., 2007, was found capable of detecting all human-pathogenic filoviruses known by 2016 (Rieger et al., 2016).

Like the assay developed in this study, the test designed by Ogawa et al., 2011 is a conventional pan-filovirus one-step RT-PCR, and its four primers (two sense and two antisense primers) target the nucleoprotein gene. The two assays differ in the number of available sequences their design had been based on and the length of their PCR product.

On the 17 November 2010, the date of the online publication by Ogawa et al., 2011, only 45 sequences of the nucleoprotein (NP) coding region of filoviruses were deposited in the GenBank database. In contrast, at the time of the primer design (17 April 2015) of the one-step pan-filovirus RT-PCR assay presented in this study 223 NP nucleotide sequences were available in GenBank. Mainly due to a large number of new sequences determined during the EVD epidemic in West Africa in 2014 – 2016 even almost ten times more NP sequences (2,219) could be found in the GenBank database for the multiple sequence alignment analysis of the designed universal consensus primer pairs (Table 2 and 3).

Specific amplification of the targeted NP sequence in a filovirus genome yields a PCR product in the assay of this study and the test developed by Ogawa et al., 2011 of 317 bp and 594 bp, respectively. The advantage of a longer amplicon (594 bp) produced by the Ogawa et al., 2011 test for species identification and virus discovery might be outweighed by the suitability of a shorter PCR product (317 bp), gained in the assay presented in this study, for further development of a dye-based real-time RT-qPCR. First applications of the assay as dye-based one-step real-time RT-qPCR, using the same universal consensus primer set, reaction and cycling conditions as designed and optimized for the conventional RT-PCR test, showed promising outcomes in increased sensitivity, real-time visualization of results, and rapid verification of specific amplicons by high resolution melt analysis (Kopp, 2015, unpublished).

Direct comparison of these previously developed RT-(q)PCR tests and the new pan-filovirus RT-PCR assay concerning sensitivity and specificity has not been possible. Templates of different origins, such as *in vitro* transcribed RNA, total RNA extracts of infected cell cultures, blood or tissues of animal models or human patients, were used in evaluating these assays. In addition, the sensitivity was measured in different units, such as RNA copies per assay or plaque- or focus-forming units in infected cell cultures. Furthermore, various sets of pathogens relevant for differential diagnosis of viral hemorrhagic fevers were employed in determining the specificity of each assay.

However, it would be beneficial to have at least two RT-PCR assays with high sensitivity and specificity each targeting a different gene of filovirus genomes. The gained complementary PCR products would allow the first confirmation of positive detection results already during the screening process of large ecological sample sets with RT-PCR assays before more time-consuming and expensive follow-up sequencing. Therefore, a systematic evaluation procedure of all currently available pan-filovirus RT-PCR assays using a standard set of specific and unspecific templates ranging from *in vitro* transcribed RNA to samples of infected animals and human patients as well as a consistent method of sensitivity measurement seems essential.

In this study, the optimization and proof-of-principle of the developed one-step pan-filovirus RT-PCR assay were conducted under sets of standardized conditions in the laboratory, which could be controlled one-by-one. However, the real-life robustness, applicability, and accuracy of this assay will have to be evaluated in the field. When applied to large scale ecological investigations into the range of potential natural hosts of filoviruses, it could be assumed that not all factors, such as source, quality, acquisition, handling, transport, and storage of the samples can be controlled at all times, especially in low resource settings of remote regions. The laboratory facilities in which the RNA extraction and RT-PCR will be performed might be of sub-standard and lack essential infrastructure and must cope with frequent power dips and error sources, such as cross-contamination by previous experiments. The actual feasibility, practicability, and ease of usage by minimally trained staff with limited resources at hand will have to be tested under field conditions. The final purpose of this test will only be achieved when it will be applied in large-scale ecological studies, which aim at finding the natural reservoir and transmission hosts of filoviruses.

